## Supplementary material for "Fast evolutionary turnover and overlapping variances of sex-biased gene expression patterns defy a simple binary sex-classification of somatic tissues": Figure 1 Figure supplement 1

**Figure 1, Figure supplement 1.** Histograms of distributions sex-bias ratios for all mouse and human organ datasets. Only significant values are included (based on Wilcoxon test statistic). To make the histograms better comparable, we plotted symmetric values for males (blue) and females (red) derived from M/F and F/M ratios >1.25. the Values >2.0 are combined into single bins.

Mouse datasets for somatic organs
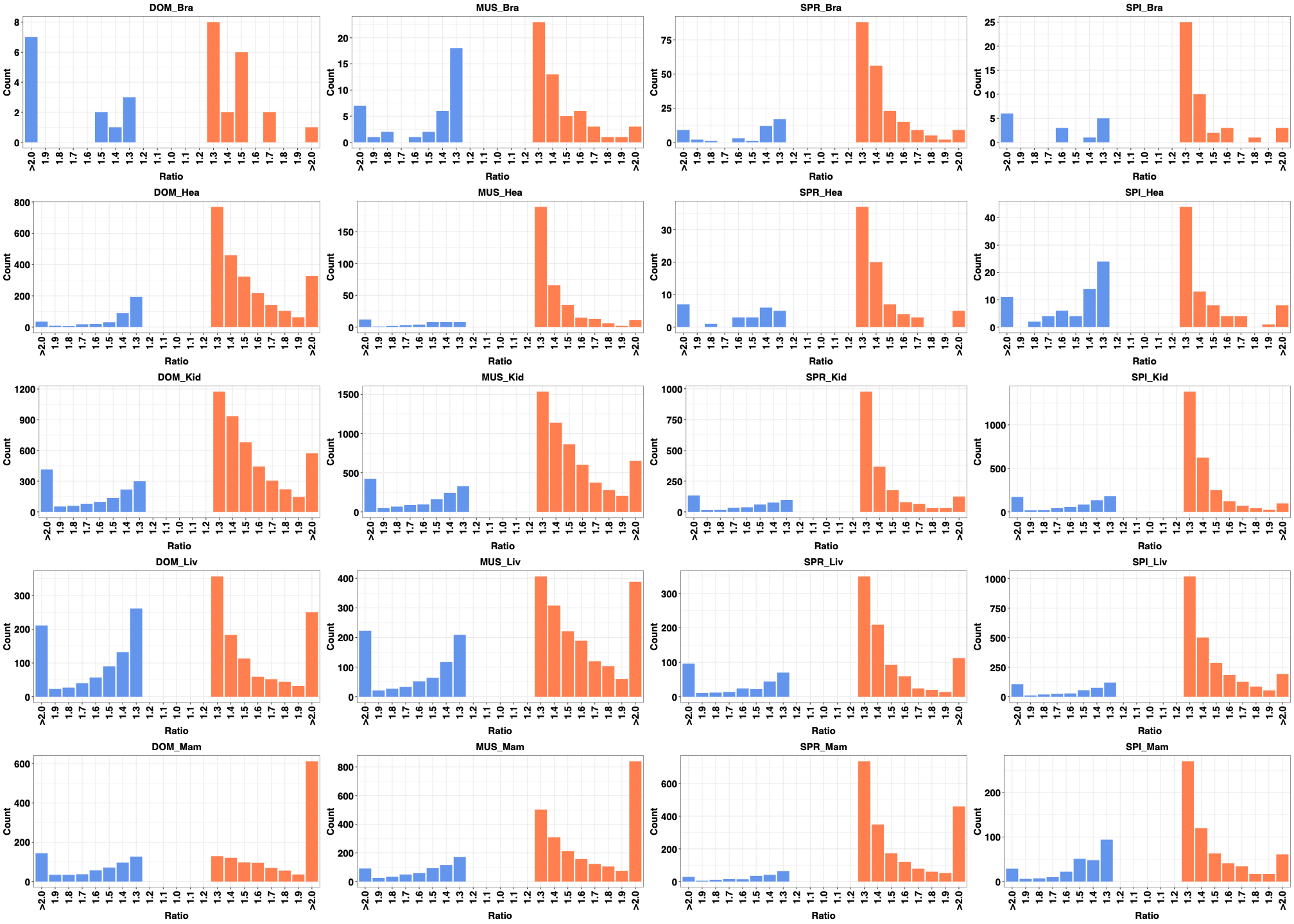

Human datasets

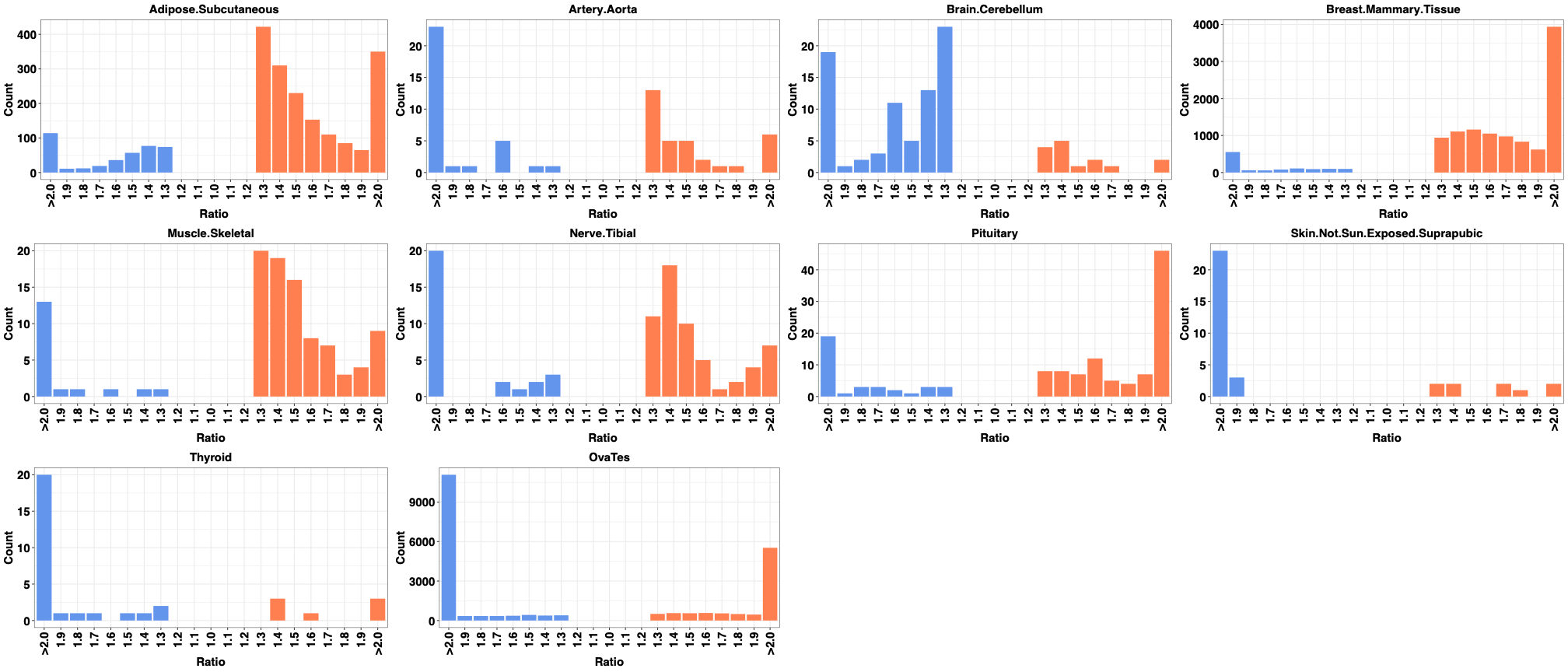
