## Supplementary figures and images for "Fast evolutionary turnover and overlapping variances of sex-biased gene expression patterns defy a simple binary sex-classification of somatic tissues"

### Figure 1 Figure supllement 2

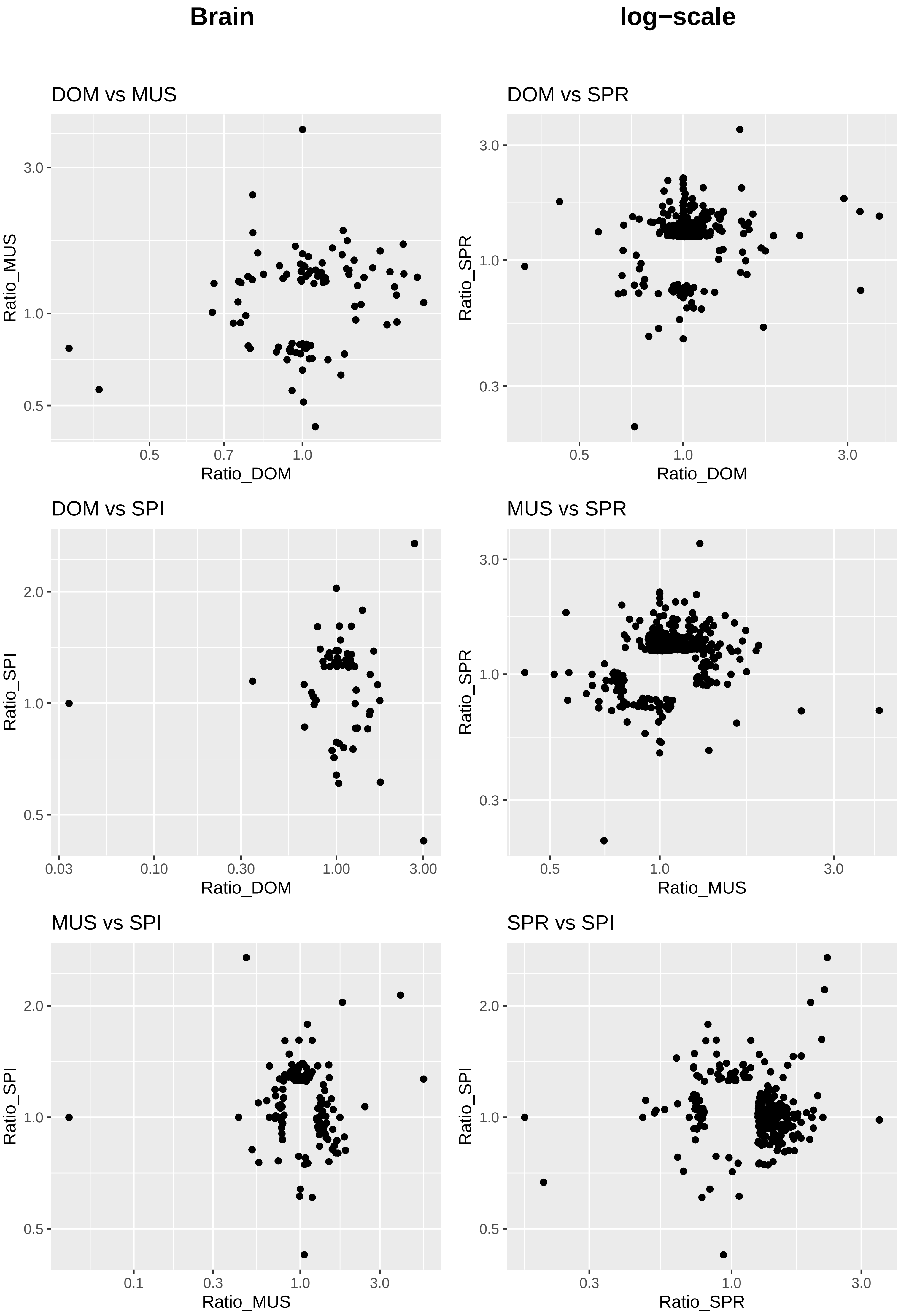

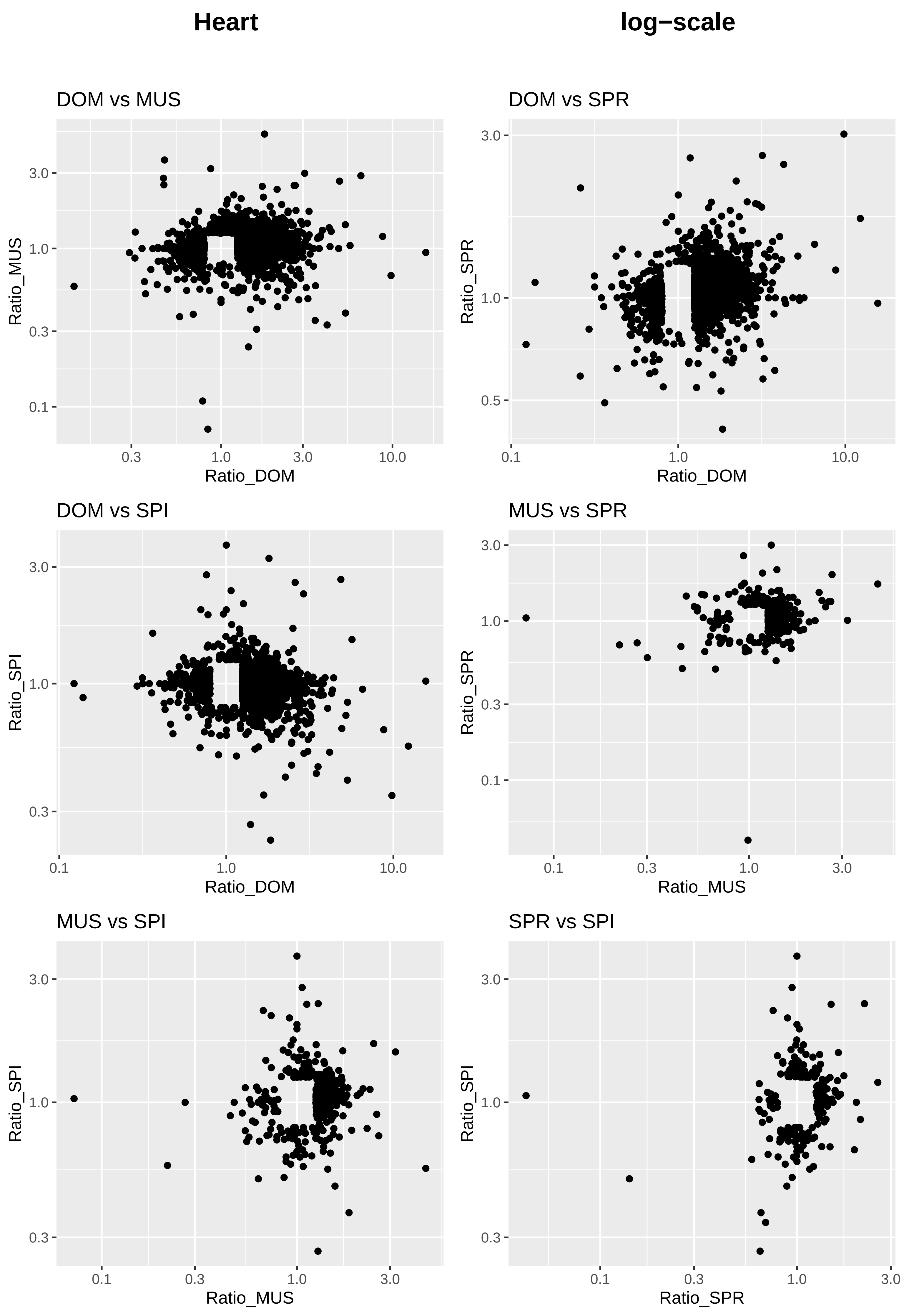

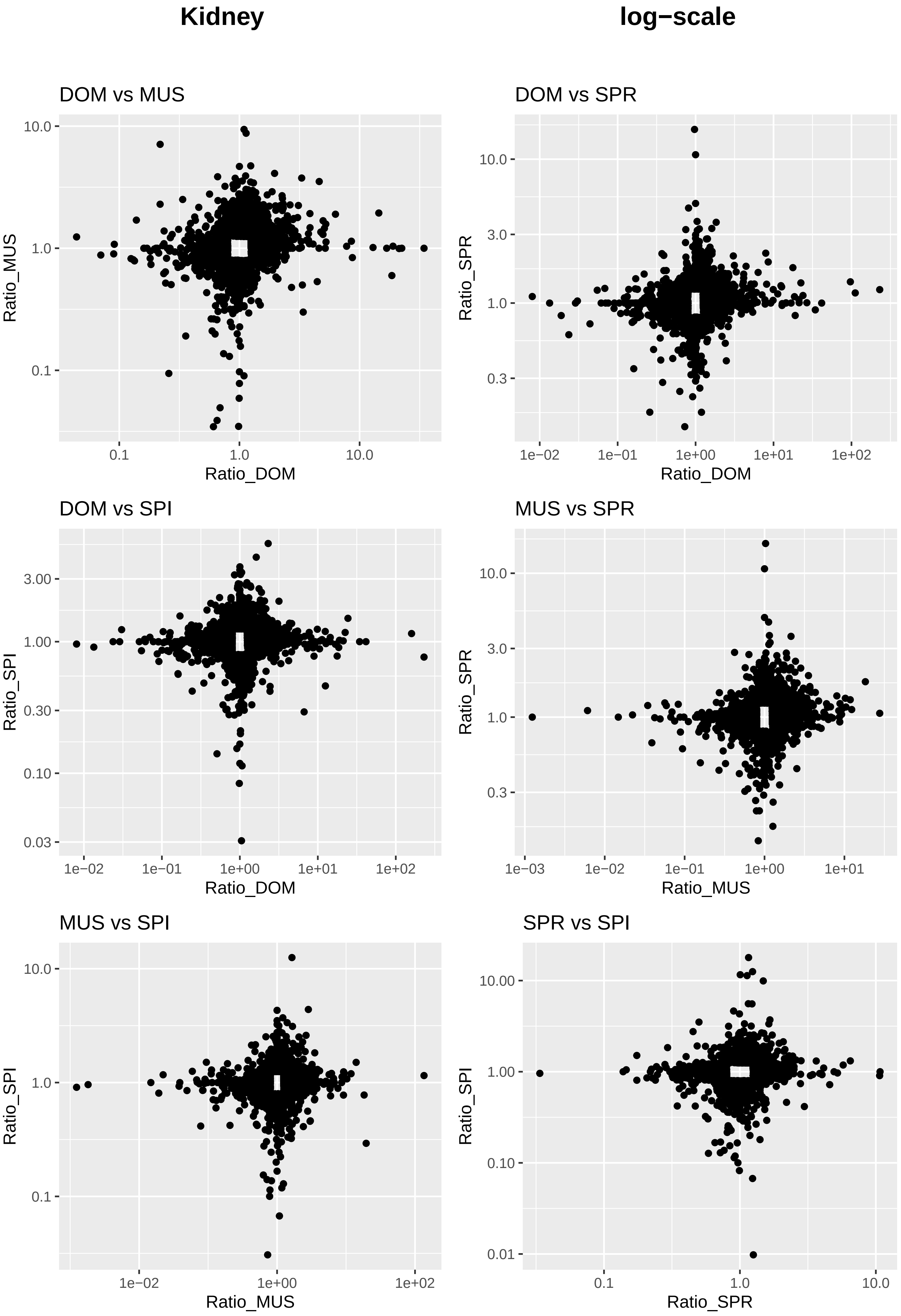

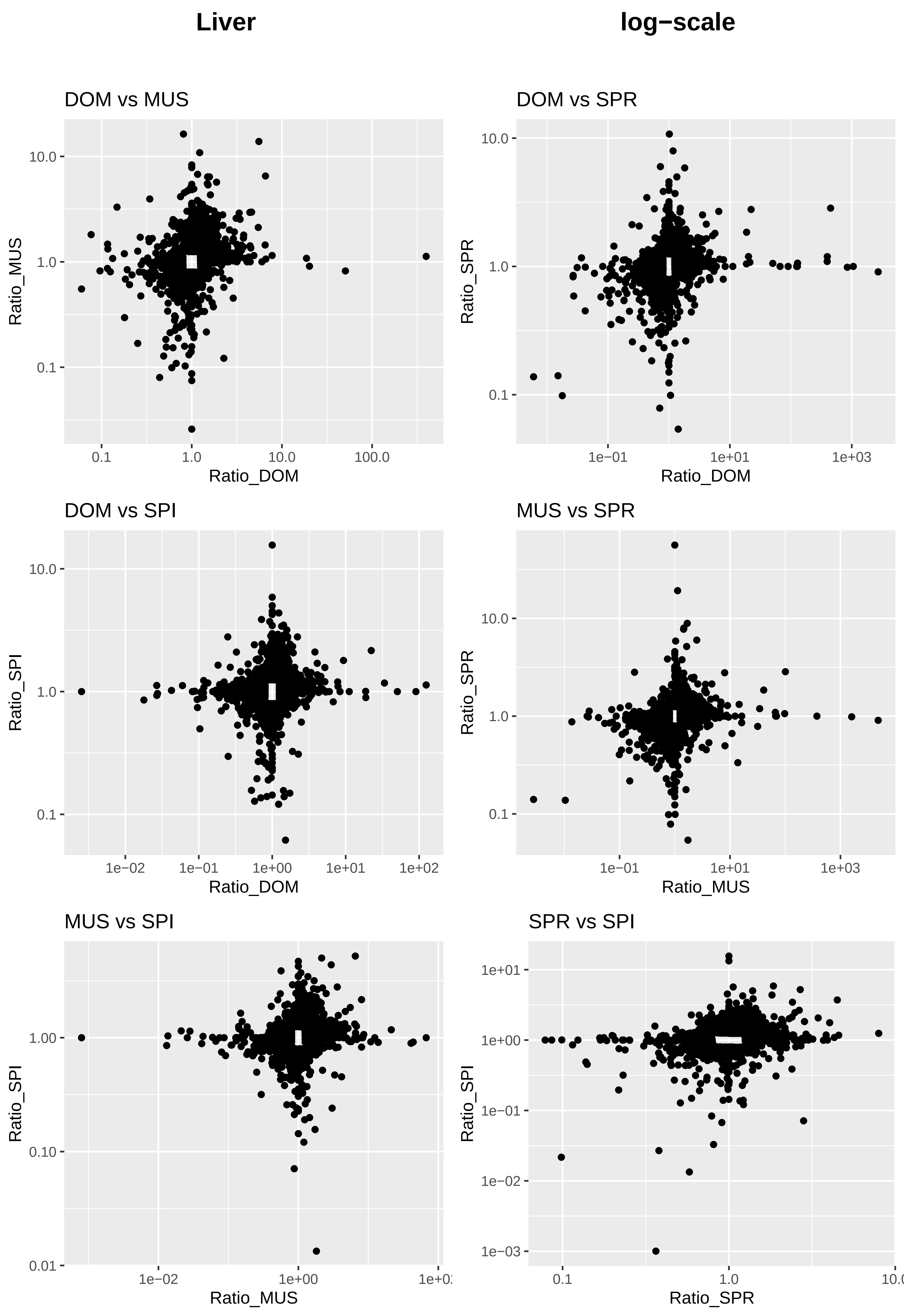

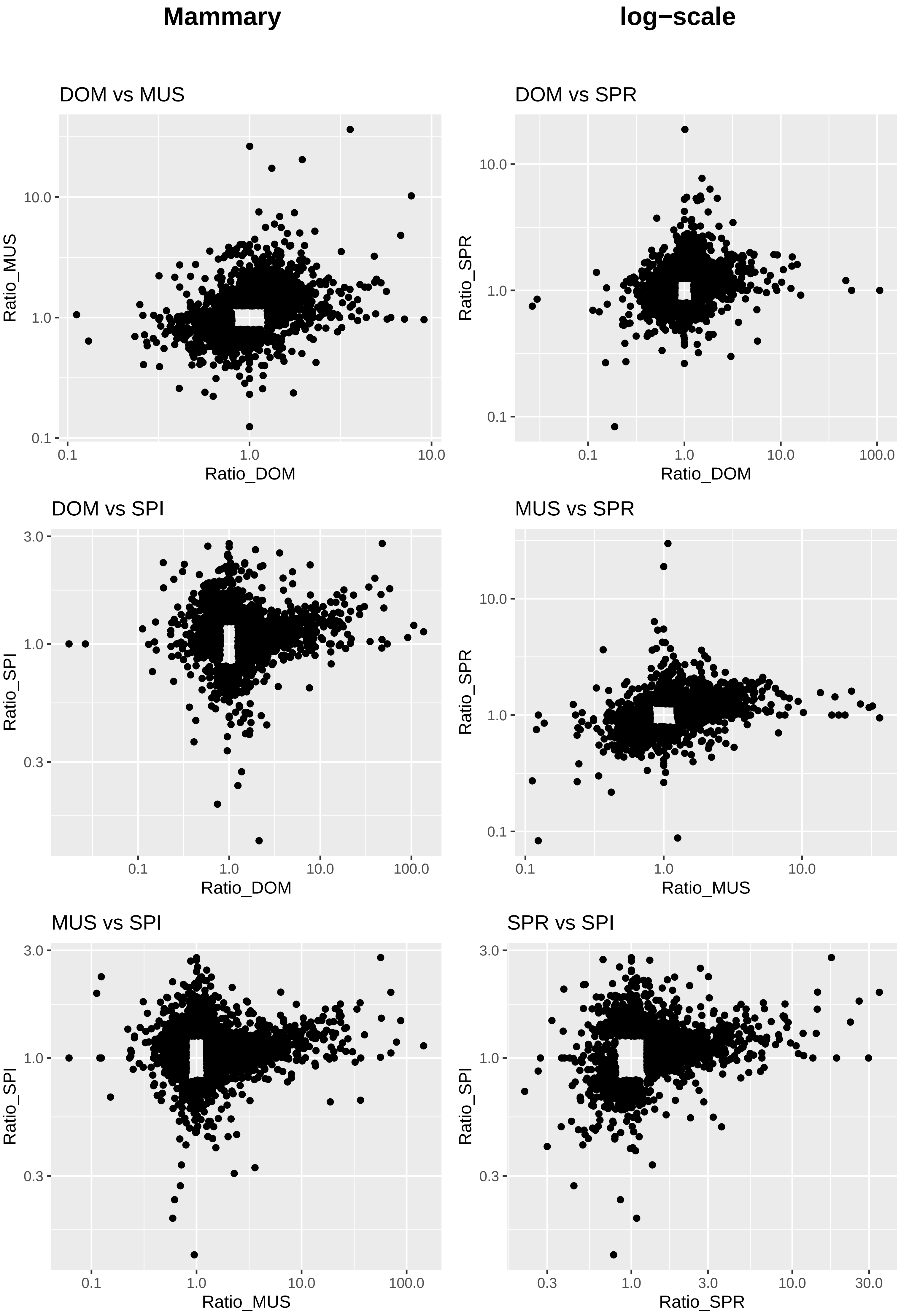

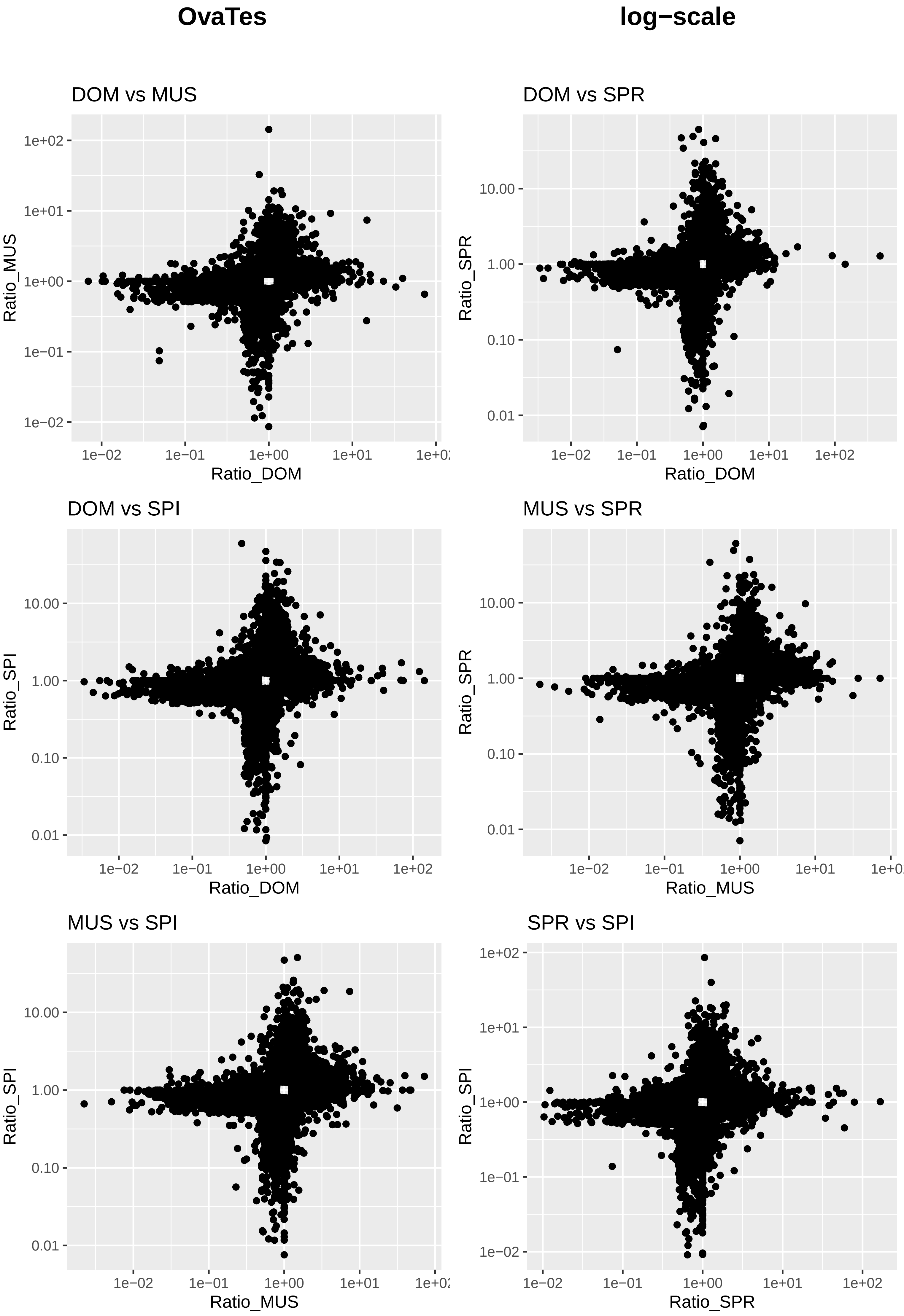

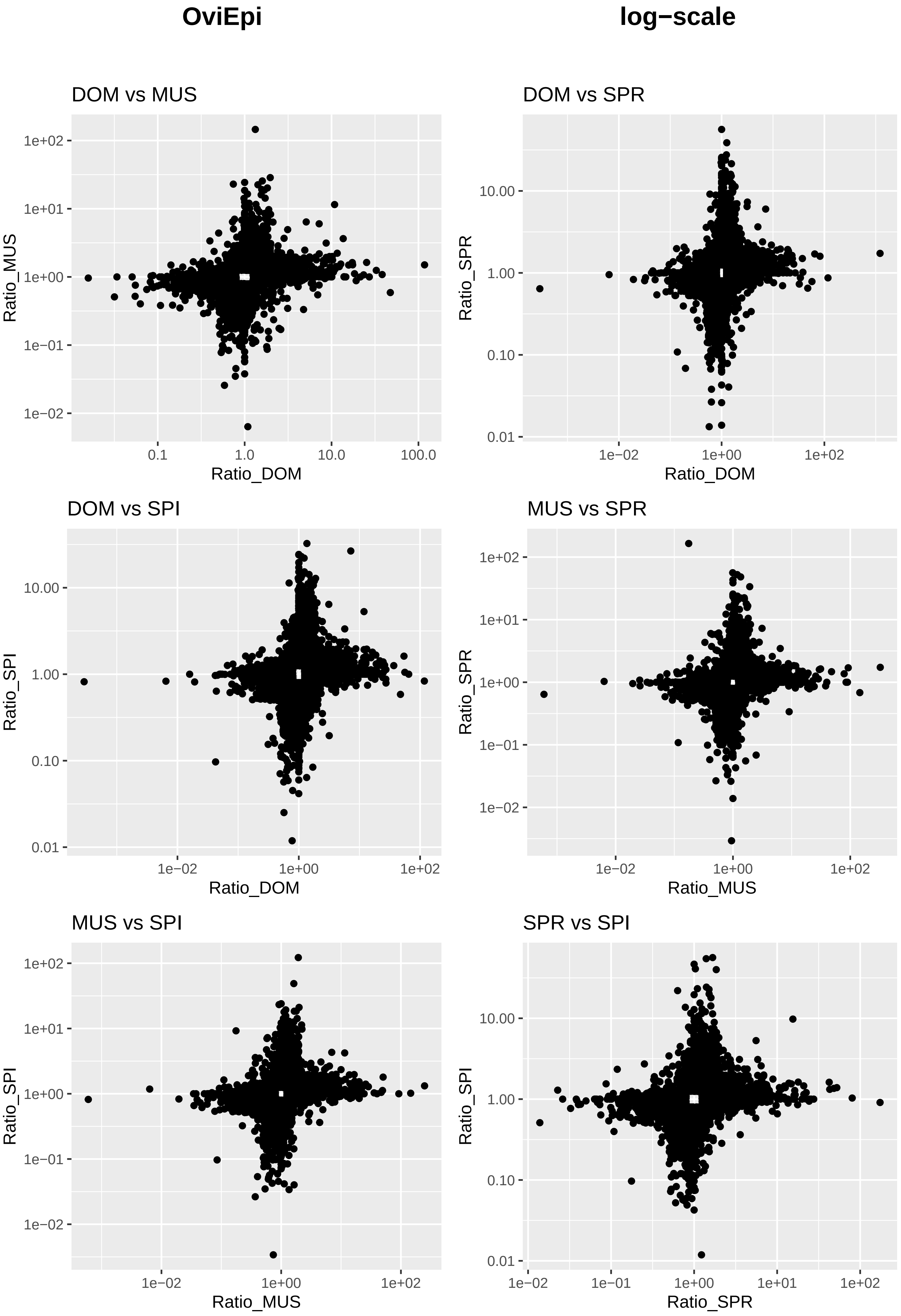

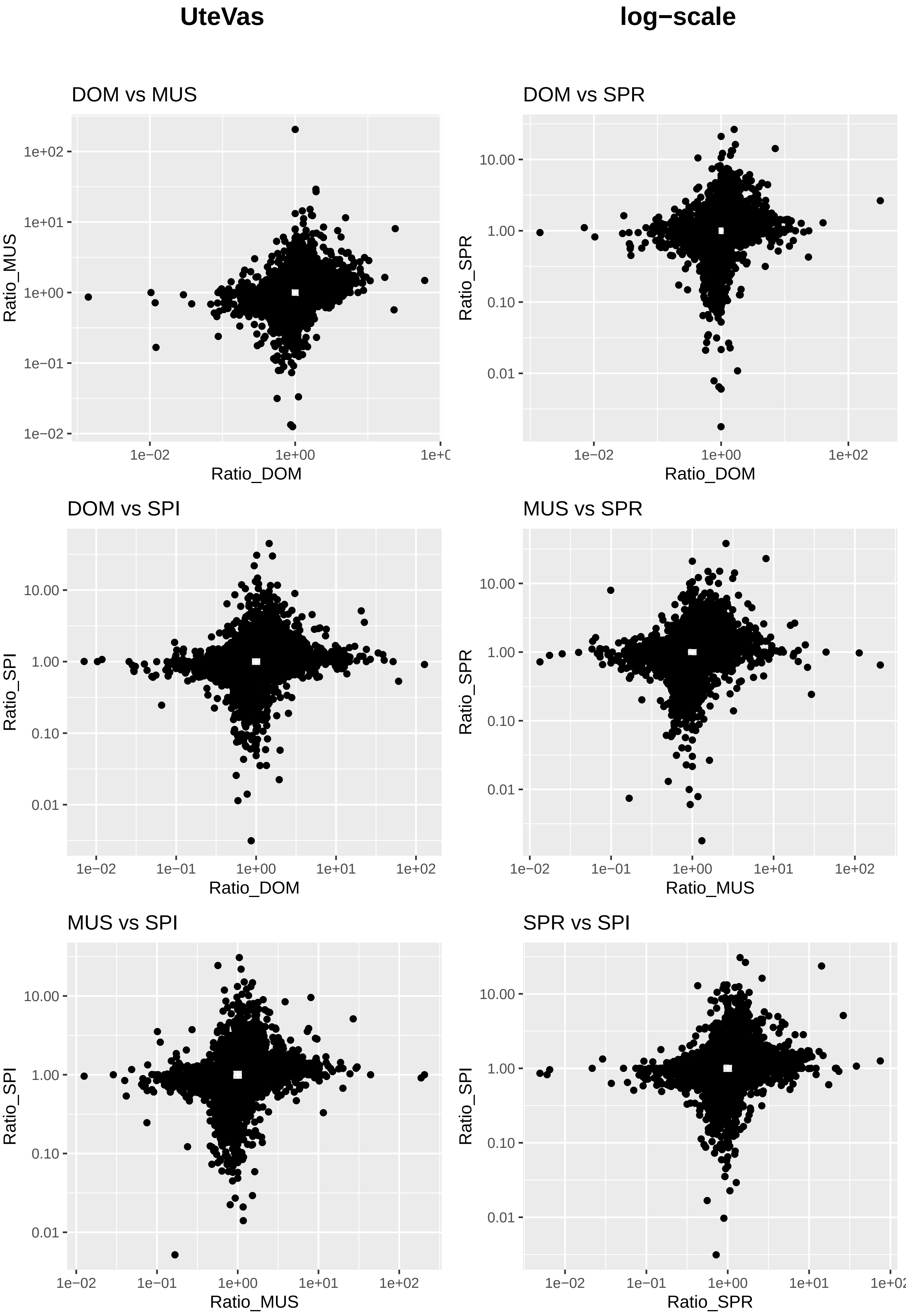
